# Using AI to Build AI: AIDO.Builder Enables Autonomous Machine Learning Model Building for Biomedicine

**DOI:** 10.64898/2026.04.20.719735

**Authors:** Han Guo, Youwei Liang, Xingyi Cheng, Caleb Ellington, Pengtao Xie, Le Song, Eric Xing

**Author notes:** Equal contribution (Han Guo, Youwei Liang).

## Abstract

Machine learning accelerates biomedical discovery, but creating effective predictive models requires specialized human expertise and demanding manual effort. Researchers must iteratively design pipelines, select architectures, and debug code. This challenge is particularly severe in biomedicine because of the heterogeneous datasets, sparse annotations, and complex evaluation protocols that are common in the domain. We present AIDO.Builder, an agentic artificial intelligence system that fully automates the entire life-cycle of biomedical model development. Provided only with a natural language task description and a target metric, AIDO.Builder autonomously constructs executable training and evaluation pipelines. The system selects suitable modeling strategies, executes experiments, and uses automated feedback-loop to iteratively revise its own code, configurations, and training procedures. It flexibly adapts to new tasks by training specialized models *de novo* or by using pretrained foundation models to build predictive models through task-appropriate adaptation. We show that across diverse biomedical benchmarks, AIDO. Builder produces highly competitive solutions against human alternatives, while eliminating the manual iteration previously required for robust model development. By automating the translation of raw data into reliable AI models, AIDO.Builder demonstrates how AI itself can be used to accelerate AI for biomedical research.

## 1 Introduction

Scientific discovery has long been driven by a systematic, iterative, often time open-end workflow invented and executed by human researchers. To uncover new knowledge, scientists would typically begin with a clear problem formulation; followed by logical experimental designs and meticulous executions; accompanied by troubleshooting of unforeseen methodological or technical failures along the way; and then analysis and theorizing of the outcomes. This flow can be loopy – by analyzing the resultant empirical evidence, researchers refine their procedures and hypotheses, and repeat the flow, driving convergence through a continuous cycle of trial, error, and adaptation. This workflow has historically been entirely manual, but in recent years, machine learning technology has been increasingly used to augment certain labor-intensive steps thereof, for example, analyzing massive amounts of complex data, such as genomic sequences or molecular structures, to predict experimental outcomes and propose new hypotheses [1, 2, 3, 4, 5, 6, 7]. However, despite the promise of machine learning as a powerful analyzer, applying these computational tools to a novel scientific problem is rarely a turnkey process, because machine learning algorithms must be custom-designed to fit the needs and properly trained to be useful. Making these algorithms work reliably still fundamentally relies on an equally tedious manual workflow [8, 9]. Rather than troubleshooting lab equipment, researchers now act as software engineers. For each new problem, they manually interpret task definitions, choose data representations, and compose complex model components. They then implement training and inference pipelines, diagnose inevitable code and integration failures, and iteratively refine their computational decisions based on experimental feedback. This modern cycle of planning, implementation, execution, and revision is working, but consumes immense time, creating a profound operational bottleneck that limits how quickly general modeling advances can be translated into reliable systems, particularly when tasks, modalities, and evaluation conventions vary widely [8, 10].

This bottleneck is particularly evident in biomedicine, a field characterized by diverse experimental tasks and inherently imperfect data [11, 12, 13]. Across biological datasets, problem formulations vary widely, annotations are frequently sparse, and standards for evaluating success remain inconsistent [14, 15, 16, 17]. Consequently, the primary barrier to scalable discovery is often the fragility of computational pipelines, where minor anomalies in data formatting or incompatible software dependencies frequently derail model development. Even when working with curated benchmarks, scientists routinely expend substantial effort simply to assemble functional code and resolve basic runtime errors. They are then forced to navigate a highly sensitive calibration process, where minor computational choices, such as how data are partitioned or how the learning objective is mathematically constrained, can drastically alter both performance and reproducibility [12, 18, 19, 20]. Because stabilizing these tools and systematically resolving code errors demands extensive manual intervention, this engineering overhead draws critical focus away from the researcher’s primary expertise: advanced experimental design, data curation, and biological interpretation.

Recent advances in large language models (LLMs) and machine learning frameworks now offer a clear path to automate this workflow [21, 22, 23, 24, 25]. LLMs can now translate plain text task descriptions into functional machine learning pipelines by directly generating code and computer commands [26, 27, 28, 29]. Furthermore, by injecting code execution outcomes into the context of LLMs, they can modify the code, correct errors, and systematically refine these pipelines [30, 31]. Alongside these algorithmic capabilities, existing machine learning frameworks provide reproducible execution environments, standardized data interfaces, and systematic experiment tracking. This combination establishes a rapid, automated feedback loop between proposing a methodological change and measuring its empirical effect. By harnessing LLMs in automating biomedical workflows, recent agentic systems have different focuses and scopes in this design space. The AI Scientist [32] targets end-to-end automation of machine learning research from ideation to manuscript generation and review. Biomni [33] instead emphasizes designing a broad biomedical action space spanning tools, databases, and protocol execution across many biomedical workflows, while BioDiscovery Agent [34] focuses on performing tool calling for genetic perturbation experiments and CRISPR-GPT [35] specializes in LLM copiloting for gene-editing experiment planning and analysis without machine learning model building. In contrast, Virtual Lab [22] is centered around AI–human research collaboration, with a human researcher and a team of LLM scientist agents that jointly handle project specification, agenda-driven scientific discussion, tool selection, and workflow design. These developments show the feasibility to build autonomous agentic systems that execute the model development cycle end-to-end (Fig. 1a), rather than acting as advisory tools that still require extensive manual implementation [28, 29].

**Fig. 1.**
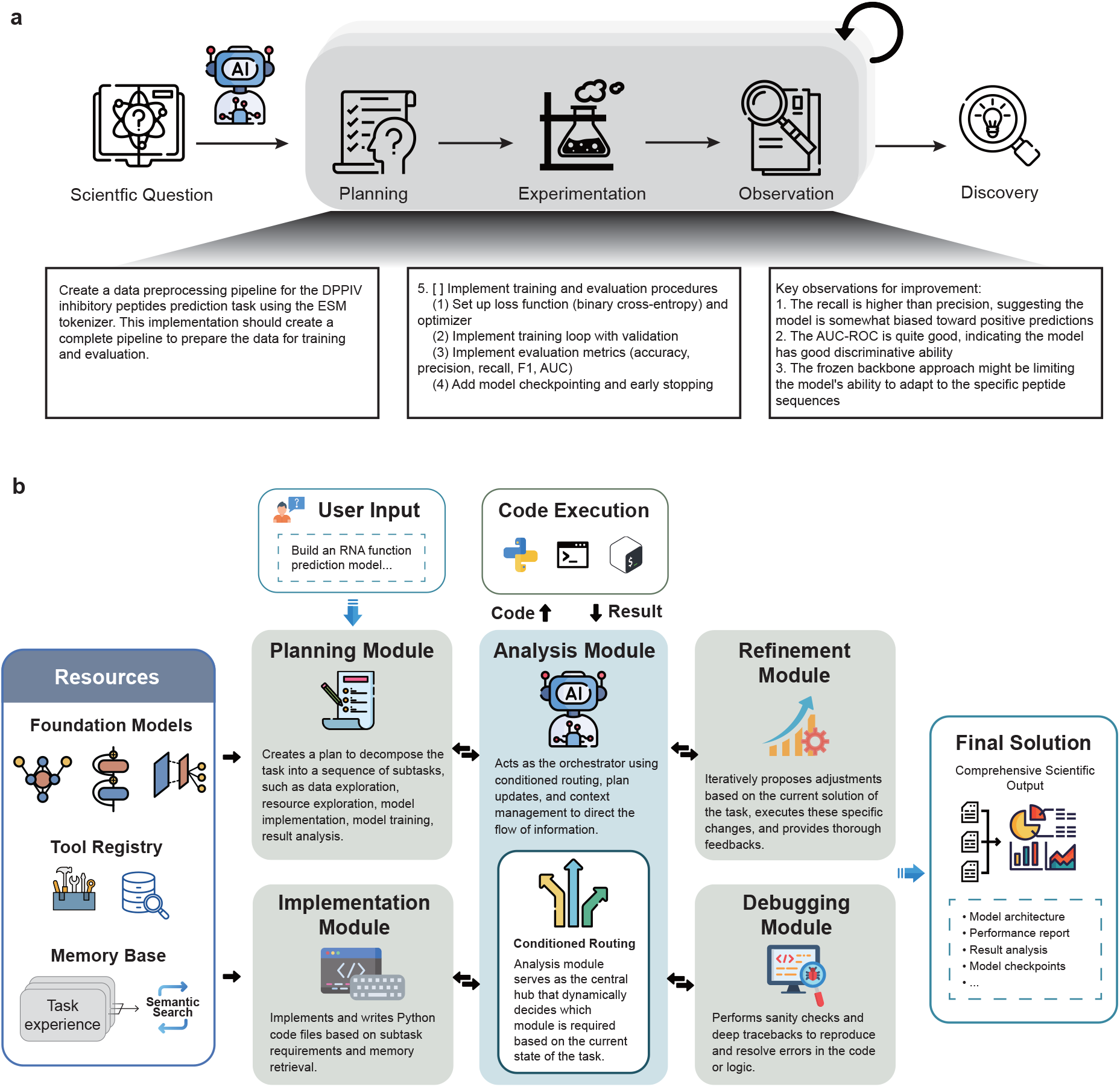
Workflow and architecture of AIDO.Builder for autonomous biomedical model development. **a**, AIDO.Builder translates a scientific question into discovery through an iterative cycle of planning and reasoning, computational experimentation, and outcome observation, providing a scalable and reproducible framework for task-specific biomedical model building. **b**, Given a biomedical task specification, AIDO.Builder approaches it using an iterative agentic workflow of planning, analysis, code implementation, debugging, and refinement, supported by resources including a suit of powerful biomedical foundation models, a tool registry, and a persistent experience memory. The planning module interprets the task, identifies constraints and available resources, decomposes the task into managable subtasks, and proposes a sequence of steps to solve them. The analysis module then acts as the central controller, maintaining task context and determining when to invoke implementation, debugging or refinement. Guided by this orchestration process, the implementation module instantiates a runnable training and evaluation pipeline, either by constructing a task-specific model from scratch or by configuring fine-tuning procedures for a pretrained foundation model. The analysis module sends the code to a virtual environment for execution, and then analyses the output. When execution reveals code or configuration failures, the debugging module diagnoses and resolves them. The refinement module subsequently revises the current solution on the basis of execution feedback and observed performance, enabling repeated cycles of improvement until the pipeline stabilizes. Underpinning this workflow is a selective, embedding-indexed memory module that accumulates structured experiences across tasks. Through similarity-based retrieval and reflection-based creation, the system autonomously transfers relevant procedural knowledge and machine learning design patterns across biologically distinct tasks.

Here we present AIDO.Builder, an agentic system that automates the full lifecycle of predictive model construction and improvement for biomedical tasks without human intervention. In contrast to The AI Scientist [32], which is positioned around scientific ideation and paper production, AIDO. Builder is designed for reliable task-level model construction and optimization. Unlike Biomni [33] and BioDiscovery Agent [34], which center tool-rich biomedical workflow execution, AIDO.Builder focuses on autonomous predictive modeling and iterative model improvement. Given a task specification and target metric, AIDO.Builder first constructs a complete, executable training and evaluation pipeline by selecting an appropriate modeling approach, exploring the provided data, and executing the resulting system in a controlled environment with both hardware and software resources (Fig. 1b). This supports tasks for which strong performance depends on well-matched, task-specific architectures trained from scratch, as well as tasks for which success depends on effective use of pretrained representations. When pretrained foundation models are beneficial, AIDO.Builder selects and configures suitable adaptation procedures to tailor them to the target prediction task, often together with task-specific prediction heads or other newly designed modules, and iteratively refines the adaptation strategy on the basis of observed outcomes. The system does not pretrain the underlying foundation models themselves; rather, it draws on pretrained biomedical foundation models, including members of the AIDO family [36, 37] as well as other task-appropriate models, alongside additional tools and resources, when constructing predictive models for downstream biomedical tasks. An integrated memory module further enables the agent to execute multi-round experimental workflows by maintaining context across iterations, allowing it to adapt its strategy dynamically, for example by shifting from biologically motivated heuristics to active-learning ensembles when experimental data are sparse. Across both regimes, AIDO.Builder uses execution feedback to guide revision, resolve code and configuration errors, and repeat this loop until performance stabilizes.

We evaluate AIDO.Builder on curated biomedical tasks that differ in task structure and their optimal modeling regimes. Across tasks, AIDO.Builder constructs viable baseline solutions from minimal task specifications and then improves them through iterative experimentation, thereby reducing the manual trial and error typically required to reach competitive performance. In settings where training from scratch is appropriate, the agent converges on strong, task-specific pipelines by selecting suitable model components and training recipes. In settings that benefit from pretrained representations, the agent adapts foundation models through automated fine-tuning and refinement of the training procedure. Together, these results show that a single agentic framework can support both task-specific model development and foundation model adaptation within a unified end-to-end workflow.

More broadly, AIDO.Builder provides a scalable route for translating general modeling advances into reliable task-level systems in biomedicine. By automating the cycle of planning and reasoning, computational experimentation, and outcome observation, it reduces the engineering burden of model development while preserving reproducibility through explicit artifacts and execution traces. This capability will become increasingly important as biomedical tasks diversify and as successful solutions depend not only on model choice, but also on disciplined system design, evaluation, and iterative optimization.

## 2 Methods

### 2.1 Agent Architecture

We formulate the autonomous research agent as a decision-making entity operating within a Partially Observable Markov Decision Process (POMDP), defined by the tuple ℳ = (*S, A, T, R*, Ω, *O*). The state space *S* represents the totality of the coding environment, including the file system, installed libraries, and external data resources. As the agent cannot observe the entire state of the external environment simultaneously, it operates on observations *O* ∈ Ω, which consist of the execution history, terminal logs, and currently read file contents. The agent’s policy *π*(*a* |*o*), parameterized by a fixed Large Language Model (LLM), maps these observations to actions *a* ∈ *A* without gradient-based training. Instead, the agent utilizes in-context learning and a structured state graph to navigate the problem space, effectively performing inference-time optimization of the solution.

#### Hierarchical planning (*v*_plan_)

The workflow begins at *v*_plan_ where the agent maps the user objective to a hierarchical task decomposition. The planner builds a structured graph of sequential subtasks and tracks progress with status indicators. This phase relies on a resource description that includes available functions, software libraries, and data lake indices. This ensures the proposed decomposition is practical within the environment.

#### Implementation (*v*_impl_)

Transitions to *v*_impl_ occur when the plan has pending tasks. In this state the agent creates atomic actions *a* ∈ *A* by writing an implementation requirement and a subtask name. The system improves generation quality by using a context selection method that selects only relevant logs and file dependencies. This reduces noise in the context window. The policy is constrained to produce concise and minimal working examples rather than complex software patterns to prioritize research progress.

#### Environment interaction (*v*_exec_)

After implementation, the state *v*_exec_ submits the actions to the environment. The agent generates a shell script limited to 300 lines to prevent complexity issues. This script runs in a secure subprocess that records standard output and runtime logs. The state produces a new observation *o*^′^ ∈ Ω that includes the execution logs and the list of file changes.

#### Code fixing (*v*_fix_)

Upon unsuccessful execution, the state *v*_fix_ performs comprehensive code debugging and fixing in existing buggy code base. The agent is instructed to investigate and explore suspicious code files before applying patch fixes. Upon fixing completion, the state *v*_fix_ quickly generates the execution script and reruns the script in a secure subprocess that records standard output and runtime logs. The state produces a new observation *o*^′^ ∈ Ω that includes the execution logs and the list of file changes, similar to *v*_exec_.

#### Semantic analysis and routing (*v*_eval_)

The cycle moves to the decision node *v*_eval_. Here the agent checks the observation *o*^′^ against the success criteria by summarizing the execution logs into a structured status report. This evaluation determines the next transition *δ*: Ω × *V* → *V*. The workflow proceeds to *v*_exec_ or *v*_impl_ if the code ran successfully. It routes to *v*_debug_ to inspect variables if runtime errors occur. The system transitions to a fixing state *v*_fix_ for automatic patching if the error is clear. If the strategy is unworkable the system returns to *v*_plan_ with feedback. The agent only reaches a terminal success state when the objective is verified.

### 2.2 Memory System

The agent maintains a persistent, embedding-indexed experience memory that accumulates structured records across experimental runs and is shared across biologically distinct tasks (Fig. 8). The memory system comprises two phases—reflection-based memory creation at the end of each task, and similarity-based memory retrieval at the beginning of each implementation step—which together enable transfer of procedural knowledge, ML pipeline design patterns, and operational lessons across tasks without explicit human curation.

#### Memory representation

Each memory item is a structured record containing four fields: (1) a natural-language task description summarizing the subtask (e.g., “Implementing an XGBoost ensemble for gene selection”), (2) a bigger context field explaining why the subtask was needed within the broader experimental workflow, (3) a best solution field describing the approach and key code patterns that worked, and (4) a lesson learned field documenting failure modes, bug fixes, and practical recommendations for future executions. Each memory item is indexed by a 1,536-dimensional embedding vector computed from the task description using the OpenAI text-embedding-3-small model [38]. The memory is persisted to a centralized database, enabling accumulation across independent agent runs.

#### Memory creation

Memory items are generated through an LLM-driven reflection process triggered upon task completion, when the agent’s workflow reaches the terminal state. The reflection proceeds in two stages. First, a “subtask identification” stage prompts a dedicated reflection LLM (using the same base model as the agent, at temperature 0.7) to analyze the summary of each step in the execution trajectory—comprising action taken, file creation and modification, and execution outcomes—and decompose it into semantically meaningful subtasks. The prompt instructs the LLM to identify “any coherent unit of work worth remembering for future tasks,” including environment setup, algorithm implementation, debugging sequences, and data processing pipelines, while deduplicating repeated occurrences and excluding trivial operations. Each identified subtask is returned as a name-description pair. Second, for each identified subtask, a memory generation stage prompts the reflection LLM to produce the structured four-field memory item. The LLM receives the summary of the execution trajectory, the focused sub-task description, and a context summary gathered by the agent’s context-generation mechanism (which selects relevant history steps and source code files pertinent to the subtask). The LLM needs to generate the requested fields, including the bigger context, best solution, and lesson learned. The textual task description is embedded via the OpenAI API. The resulting memory item is immediately appended to the persistent store and saved to disk, making it available for retrieval in subsequent runs.

#### Memory retrieval

The agent retrieves relevant prior experiences when the planner generates an implementation plan for a new subtask. At the retrieval point, the current task or subtask description is embedded using OpenAI `text-embedding-3-small`, and the memory store is searched by computing cosine similarity between the query embedding and all stored task-description embeddings. The five most similar items with similarity above a threshold of 0.8 are returned per retrieval, which ensures the retrieved tasks are relevant to the query. Retrieved memory items are formatted into a “related experiences” block that is injected into the implementation prompt provided to the coding LLM. Each retrieved experience is presented as a previous task description followed by its best solution, bigger context, and lessons learned. The LLM is instructed to refer to the related experiences but is not required to follow them, allowing it to selectively adopt relevant patterns while disregarding inapplicable content.

Critically, the retrieval mechanism operates purely on subtask level based on semantic similarity of task descriptions, which enables fine-grained memory recall and learning from details of past subtask experience. As a consequence, experiences from biologically similar tasks (e.g., a lysosomal choline recycling screen) can be retrieved when working on a different task (e.g., an interferon-gamma regulation screen) if their subtask descriptions are semantically similar (e.g., both describe “implementing a machine learning model for gene perturbation prioritization”). This design enables broad transfer of task-agnostic engineering patterns, such as environment variable configuration, gene submission retry logic, and ML pipeline architecture.

### 2.3 Iterative Refinement

#### Optimization setting and notation

To maximize the scientific efficacy of the generated models, we treat the refinement process as a search problem over the space of valid training pipelines. Formally, given an initial solution state *S*_0_ (the draft model), the module seeks a state *S*^*^ such that the performance metric Φ(*S*^*^) *>* Φ(*S*_0_), where Φ can be either single performance metric (e.g. AUC-ROC), or a set of performance metrics (e.g. accuracy, F1 score, and recall). The system actively traverses this solution space by iteratively mutating model architectures and systematically tuning hyperparameters based on the analysis of the current solution. This ensures that the agent does not settle for local optima but constructively evolves the solution to maximize its theoretical potential and empirical robustness.

#### Dependency modeling for safe mutation

A fundamental challenge in autonomously evolving complex software is maintaining structural integrity across interdependent modules. Aggressive architectural mutations—such as altering module connectivity or layer depth—can frequently disrupt dependencies in lower-level utilities, leading to cascading system failures. To mitigate this risk, our framework models the generated codebase as a Directed Acyclic Graph (DAG), *G* = (*V, E*), where *V* represents the set of code modules and *E* represents directed import dependencies. Prior to modification, the system computes a topological ordering of *G*. Modifications are strictly sequenced according to this hierarchy: files with the fewest dependencies (leaf nodes) are stabilized first. This dependency-aware scheduling ensures that foundational utilities are validated before dependent high-level modules are restructured, preventing inconsistent function signatures.

#### The refinement cycle: proposal, execution, and evaluation

The optimization process functions through a continuous cycle comprising three distinct phases:

#### Proposal phase

The agent analyzes the current solution state *S*_*t*_ to identify performance bottlenecks (e.g., overfitting or slow convergence). It then formulates a mutation strategy Δ, synthesizing code that targets either model architectural change or hyperparameter tuning (adjustments to the training pipeline).

#### Execution phase with code fixing

Following proposed improvement implementation, the agent initiates the training pipeline. This phase includes an stand-alone debugging mechanism. If the proposed mutation Δ introduce syntax errors or runtime exceptions, the system captures the error trace and triggers a sub-loop to diagnose and patch the implementation in real-time. This ensures that architectural exploration is not halted by trivial implementation bugs.

#### Evaluation phase

The cycle concludes with an analytical assessment. Rather than relying solely on scalar metrics, the system reasons over the training trajectory (analyzing stability and generalization gaps) to determine the validity of the new solution *S*_*t*+1_. Based on this analysis, the system accepts *S*_*t*+1_ only if it represents a robust improvement over the baseline *S*_*t*_.

## 3 Results

### 3.1 Overview of AIDO.Builder workflow

To address diverse biomedical machine learning challenges, we designed AIDO.Builder to operate as an iterative, agentic workflow comprising planning, analysis, implementation, debugging, and refinement. Upon receiving a task specification, the system establishes an initial strategy by evaluating constraints and available resources. A central analysis module orchestrates the workflow, directing the code module to instantiate a functional training and evaluation pipeline—either by constructing models de novo or by configuring fine-tuning procedures for pretrained foundation models. Execution errors are autonomously diagnosed and corrected by a dedicated debugging module, while a refinement module continuously optimizes the solution based on empirical performance feedback until the pipeline stabilizes. For open-ended biological discovery tasks, this core workflow is augmented by a selective, embedding-indexed memory module that couples task-end reflection with similarity-based retrieval to autonomously transfer procedural knowledge and pipeline design patterns across distinct biological contexts.

### 3.2 Autonomous formulation of task-specific modeling pipelines

Translating natural language task specifications into functional machine learning systems requires the autonomous design of task-specific computational structure. We evaluated the agent’s structural formulation capability using the Histopathologic Cancer Detection task [39], which requires binary classification of lymph node image patches according to whether the image contains metastatic tissue (Fig. 2a). Rather than generating a monolithic script, the agent organized the problem into a modular pipeline architecture spanning dataset inspection, preprocessing and augmentation, predefined data split handling, custom dataset construction, model definition, optimization, and validation. This decomposition constituted a coherent structural design matched to the task. Specifically, image patches were processed through a transfer-learning classifier based on a pretrained ResNet18 backbone; the prediction layer was adapted to binary classification; and model selection was aligned with the external evaluation criterion through AUROC-based validation. These results indicate that the agent was able to infer an appropriate computational scaffold directly from the natural language task specification, cleanly separating data processing, representation learning, optimization, evaluation, and output generation into interoperable modules. Comparative evaluation against a baseline agent, Biomni [33], demonstrated the superior discriminative capacity of this autonomously generated pipeline. Specifically, AIDO.Builder achieved an Area Under the Receiver Operating Characteristic (AUROC) of 0.987, outperforming Biomni’s AUROC of 0.943 (Fig. 2c). Furthermore, analysis of the prediction score distributions by true class revealed that AIDO.Builder produced a substantially sharper separation between the positive and negative classes compared to the baseline (Fig. 2d).

**Fig. 2.**
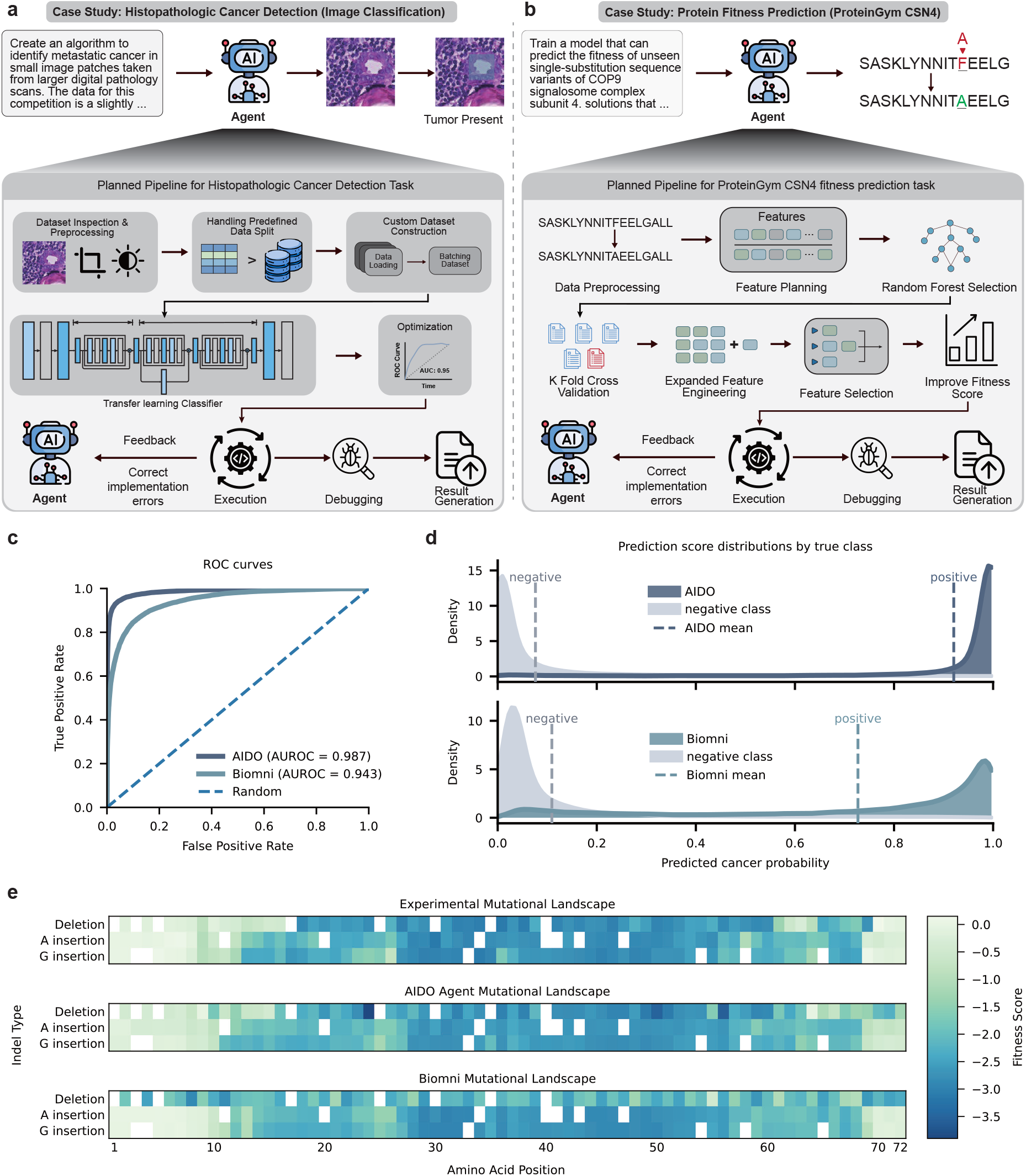
AIDO.Builder autonomously formulates task-specific modeling pipelines. **a**, Planned pipeline for the Histopathologic Cancer Detection task. From a natural-language task specification, the agent constructs a modular image-classification workflow spanning dataset inspection, preprocessing and augmentation, use of the predefined data split, custom dataset construction, transfer-learning model definition, optimization, validation and submission generation, with iterative execution, debugging and refinement. **b**, Planned pipeline for the ProteinGym CSN4 fitness prediction task. From the task description, the agent assembles a regression workflow for protein variant-effect prediction, including preprocessing, feature planning, random-forest model selection, strict five-fold cross-validation, representation expansion, feature selection and iterative improvement guided by validation feedback. **c**, ROC curves for the histopathologic classification task, showing stronger discrimination by AIDO.Builder than by the baseline method. **d**, Distributions of predicted cancer probabilities stratified by true class for AIDO.Builder and the baseline method, showing improved separation of negative and positive samples by AIDO.Builder ‘s the autonomously designed pipeline. **e**, Experimental and predicted protein fitness landscapes for indel variants across sequence positions, comparing ground-truth measurements with predictions from AIDO. Builder and the baseline method; AIDO.Builder more closely recapitulates the experimental fitness structure.

In addition to structural design, we evaluated the agent’s representation planning and dynamic optimization on the ProteinGym CSN4 [40] task (Fig. 2b). This supervised regression problem requires predicting the structural stability impacts of 196 variants (comprising 132 substitutions and 64 single-residue deletions) under a strict five-fold cross-validation protocol. The agent inferred the mutational regime and autonomously constructed a baseline feature set encoding amino acid composition, mutation count, positional information, deletion status, and relative property changes. Paired with a random forest regressor evaluated across the prescribed folds, this initial configuration yielded a Spearman correlation of 0.8741.

Following the baseline implementation, the agent executed an iterative refinement plan, utilizing empirical validation feedback to guide subsequent modeling choices. An initial attempt at nested hyperparameter optimization decreased the validation correlation to 0.7331. The agent registered this performance regression, discarded the hyperparameter configuration, and shifted its strategy to expand the representation space. It engineered 504 candidate features, incorporating *k*-mer composition, local sequence-context descriptors, and secondary-structure propensities. To mitigate dimensionality constraints, the agent autonomously applied feature importance selection, reducing the feature space to 40 informative variables. This targeted representation engineering, constrained by the task’s cross-validation protocol, recovered and improved predictive performance to a final Spearman correlation of 0.8820. These results demonstrate that autonomous modeling workflows depend on both modular architecture generation and the capacity to dynamically adjust feature representations in response to validation metrics. Furthermore, a comparative analysis of the predicted mutational landscapes (Fig. 2e) reveals that the AIDO.Builder’s generated landscape more accurately captures the experimental ground truth. While the Biomni baseline yields a comparatively smoothed and less detailed fitness map, AIDO. Builder effectively resolves the localized fitness score penalties across various indel types and amino acid positions.

### 3.3 Systematic error resolution and pipeline stabilization

The automated construction of machine learning pipelines frequently results in runtime errors, including data formatting anomalies, integration failures, and dependency conflicts. We evaluated the agent’s capacity to autonomously diagnose and resolve these errors using the PolarisHub CYP2D6 substrate identification [41] task, a binary classification problem predicting metabolic turnover from SMILES [42] strings. The agent built a molecular property prediction pipeline using 48 physicochemical and topological descriptors, explicitly handling class imbalance and using AUPRC for model selection. During initial execution, the feature extraction step yielded an empty matrix, causing a preprocessing error (Fig. 3a). The agent analyzed the traceback and corrected the computation by removing incompatible topological features and modifying the SMILES parsing logic to ensure consistent feature coverage.

**Fig. 3.**
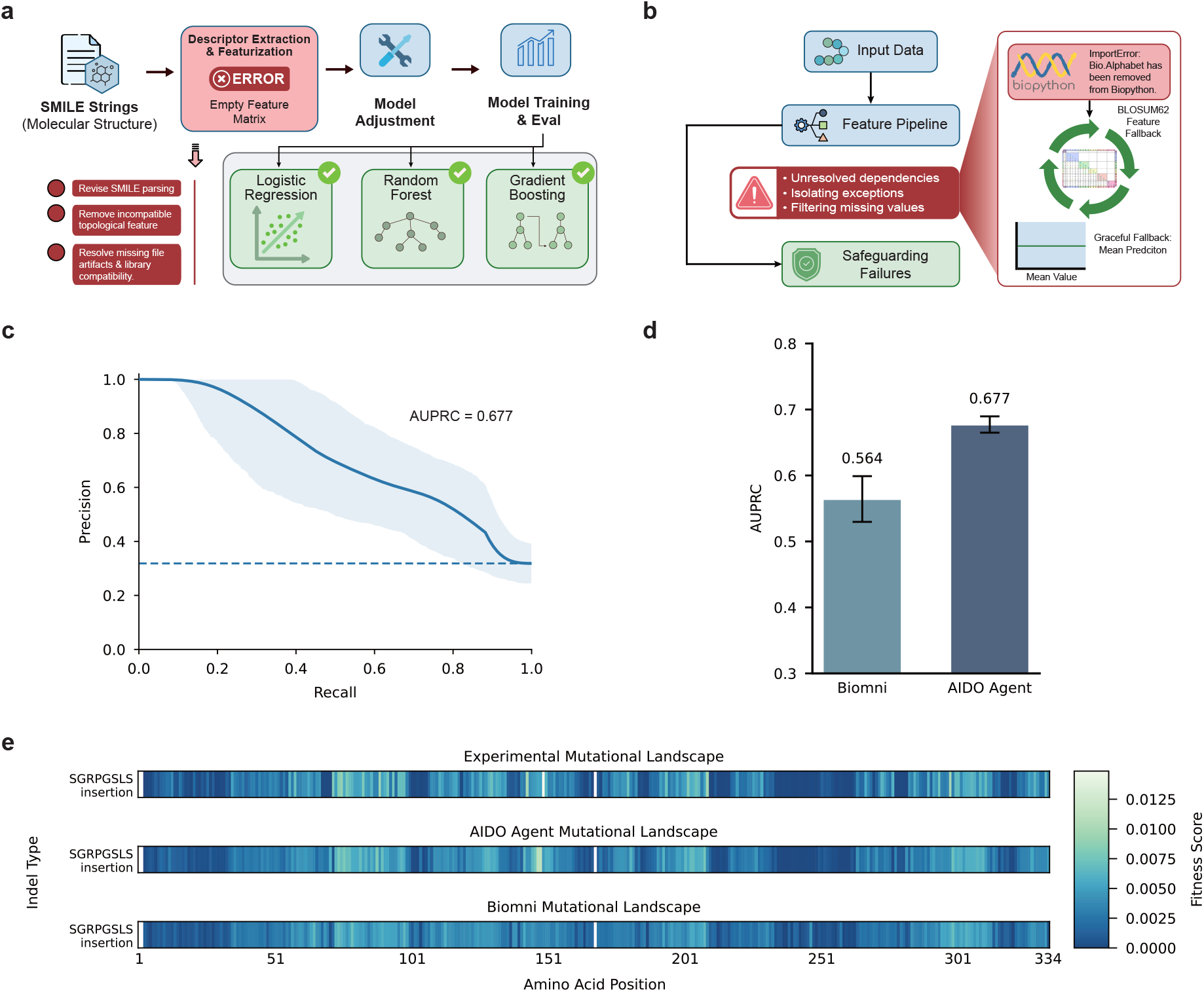
AIDO.Builder systematically resolves errors and stabilizes pipeline. **a**, Autonomous debugging on the PolarisHub CYP2D6 substrate identification task. The agent repairs multiple execution failures, including empty feature extraction, data-flow mismatches and library incompatibilities, to restore a complete molecular property prediction pipeline from SMILES strings. **b**, Robust execution on the ProteinGym Q8EG35 insertion/deletion fitness prediction task across three independent runs, showing adaptive stabilization through defensive feature engineering, exception isolation, dependency substitution, tensor reshaping and fallback execution when full recovery is not possible. **c**, Precision–recall curves for the CYP2D6 task. **d**, AUPRC comparison between AIDO. Builder and the Biomni baseline on CYP2D6 substrate identification. **e**, Q8EG35 indel fitness predictions, comparing predicted and experimental fitness scores.

The agent also systematically resolved subsequent integration errors, including a missing file artifact and a neural network library compatibility issue. It matched the data saving format with the downstream loader, verified array shapes before model training, and adjusted the multilayer perceptron configuration. These corrections enabled all candidate models, including logistic regression, random forests, and gradient boosting, to execute successfully. Following the stabilization of the computational pipeline, we evaluated the agent’s predictive performance on this imbalanced classification task. Quantitative analysis showed that AIDO.Builder achieved an Area Under the Precision-Recall Curve (AUPRC) of 0.677 (Fig. 3c), outperforming the Biomni baseline (AUPRC = 0.564, Fig. 3d).

To rigorously evaluate system robustness, we assessed the autonomous agent across three independent trials using the ProteinGym Q8EG35 insertion/deletion (indel) fitness prediction task [40]. This task requires predicting continuous fitness landscapes from mutant sequences under a strict five-fold cross-validation protocol, quantified via Spearman’s rank correlation coefficient (*ρ*) [43]. Across these trials, the agent demonstrated highly adaptive resilience under escalating workflow complexity. In the optimal trial (Run 1; *ρ* = 0.7386), the agent prioritized execution stability by structurally decoupling cross-validation from candidate model evaluation. Furthermore, it autonomously engineered a parsimonious feature pipeline, extracting 34 sequence-derived features per variant, and implemented defensive programming mechanisms. These included bypassing unresolved third-party dependencies, isolating model-specific evaluation exceptions to prevent global script termination, and stabilizing metric computation through missing-value filtration. Collectively, these architectural decisions illustrate the agent’s capacity to preserve end-to-end execution and evaluation fidelity despite component-level instability.

The second trial (Run 2) tested the agent’s stability under elevated feature-engineering demands. Here, the agent autonomously constructed a highly dimensional representation of the inserted peptides, integrating amino acid composition, physicochemical properties, localized contextual metrics, and secondary structure propensity to yield a 331 × 62 feature matrix. Despite this increased computational complexity, the agent successfully conserved the requisite cross-validation protocol, executed fold-specific preprocessing, and systematically evaluated an ensemble of candidate regressors. Although this elaborate workflow did not surpass the predictive accuracy of Run 1, its successful completion demonstrates the system’s ability to maintain operational stability and programmatic integrity while navigating broader modeling spaces.

Crucially, the third trial (Run 3) revealed the agent’s capacity to mitigate cascading pipeline failures. Upon encountering a critical dependency error due to an unavailable BioPython module (Fig. 3b), the agent dynamically repaired the script by substituting an internal BLOSUM62 [44] representation, thereby averting an import-time failure. Subsequently, a deeper architectural mismatch emerged when upstream feature branches generated three-dimensional tensors that were incompatible with a downstream two-dimensional normalization requirement. The agent executed targeted interventions, including automated tensor reshaping and feature processing adjustments, to resolve this dimensional inconsistency. When these localized repairs proved insufficient to fully stabilize the primary modeling trajectory, the agent successfully contained the failure cascade. It autonomously pivoted to a conservative fallback protocol, assigning a constant mean fitness estimate to all variants. This capacity for fail-safe execution highlights a fundamental robustness property of the autonomous system: the capability to autonomously diagnose exceptions, implement localized computational repairs, and ensure complete pipeline execution even when full recovery is intractable. Beyond operational robustness, evaluation of the predicted mutational landscapes from the optimal trial (Fig. 3e) demonstrates that AIDO.Builder more closely approximates the experimental ground truth. Compared to the generalized fitness profile produced by the Biomni baseline, the AIDO.Builder pipeline yields enhanced resolution of localized, sequence-specific fitness variations.

### 3.4 Metric-driven model selection and construction

Optimizing predictive models for biological data requires aligning the training process with the final evaluation metric and combining different models to prevent overfitting. We evaluated the agent’s ability to perform metric-driven model selection and ensemble construction using the OpenProblems Spatially Variable Genes task [45, 46]. This task requires ranking genes based on the spatial structure of their expression across tissue coordinates, using simulated ground-truth data that mixes spatial and non-spatial components. Performance is measured by the rank correlation between predicted and true gene variability scores (Fig. 4a). We compared the agent against a general biomedical baseline that uses a fixed set of global autocorrelation statistics, such as Moran’s *I* [47] and Geary’s *C* [48], and selects a single model based on cross-validation.

**Fig. 4.**
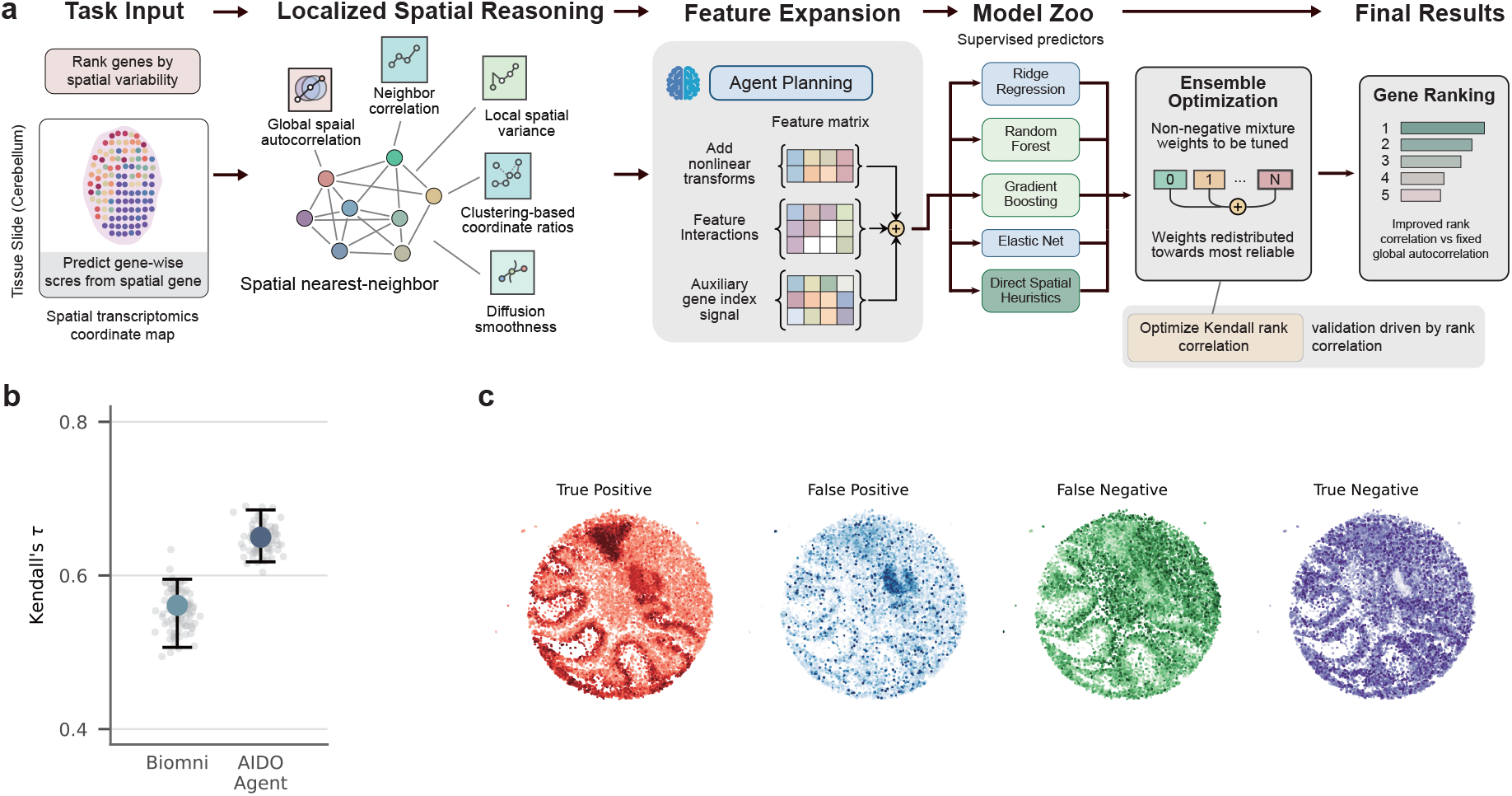
AIDO.Builder selects and constructs model with metric-driven approach. **a**, Pipeline for the Open-Problems Spatially Variable Genes task. From the task specification, AIDO.Builder constructs a modular workflow that combines localized spatial reasoning, feature expansion, supervised model selection and ensemble optimization. The agent builds a spatial nearest-neighbor graph over tissue coordinates, extracts gene-level descriptors including global autocorrelation, neighbor correlation, local spatial variance, clustering-based coordinate ratios and diffusion smoothness, expands the representation space through nonlinear transformations and feature interactions, and optimizes a composite predictor to maximize Kendall’s rank correlation on the calibration set. **b**, Comparison of Kendall’s *τ* on the benchmark, showing improved rank correlation for AIDO.Builder relative to the Biomni baseline. **c**, AIDO.Builder predicted spatial expression patterns in mouse cerebellum Slide-seqV2 grouped by prediction outcome. True-positive genes exhibit clear anatomically structured expression, whereas true-negative genes remain comparatively spatially uniform. Dots denote individual spatial beads, colour indicates mean normalized expression, and the grey background outlines the tissue.

In contrast, the agent built a pipeline focused on local spatial consistency rather than relying solely on global summaries. It constructed a spatial nearest-neighbor graph connecting adjacent tissue spots, specifically using a six-neighbor structure. From this graph, the agent extracted a comprehensive set of gene-level features. These included global autocorrelation statistics, direct neighbor correlation measures, local spatial variance, clustering-based coordinate ratios, and a diffusion-based smoothness score. Furthermore, the agent autonomously executed a second refinement pass that expanded the feature space by introducing nonlinear transformations and explicit feature interactions. It also incorporated the gene index as an auxiliary variable to capture underlying ordering signals. Recognizing that the benchmark evaluates relative gene ranking rather than absolute prediction error, the agent aligned its validation strategy by explicitly optimizing for Kendall’s rank correlation [49] on the validation dataset.

Crucially, the agent did not simply select the single best-performing model. It trained a diverse set of supervised models, including ridge regression, random forests, gradient boosting, and elastic net variants. These models were evaluated alongside direct spatial heuristics. Treating these predictors as complementary rather than interchangeable, the agent created a composite ensemble. It optimized non-negative mixture weights to directly maximize the validation correlation. This refinement increased the internal validation correlation from 0.6946 under equal weighting to 0.6956. The optimization process autonomously concentrated the ensemble mass on the most reliable components, assigning specific fractional weights to the internal ensemble predictor, ridge regression, and random forests, while maintaining smaller contributions from diffusion and gradient boosting models.

During the execution of this pipeline, the agent maintained a modular structure that separated data loading, schema verification, feature extraction, and solution assembly. This separation allowed it to enforce shape and range checks on the final gene-wise score outputs and to systematically diagnose runtime errors. It rectified erroneous assumptions about input file paths, adapted its data loading logic when intermediate feature files were missing, resolved mismatches in file and label-source assumptions during data ingestion, and addressed a mathematical edge case in the weight optimization step that returned an empty value. Finally, because small calibration datasets are susceptible to statistical anomalies, the agent autonomously implemented a feature sanity check. During validation, it identified and flagged an abnormally high-correlation variable as potential data leakage, shifting the ensemble weights toward valid spatial descriptors. By combining localized feature engineering, metric-aligned ensembling, and targeted error resolution, the agent achieved a final test rank correlation of 0.6411, compared to 0.4983 for the baseline.

While the metric-driven pipeline of the agent substantially improved overall quantitative performance, qualitative evaluation of the spatial expression profiles (Fig. 4c) reveals universally persistent challenges inherent to modeling highly sparse spatial transcriptomics data. Specifically, both the agent and current baseline methodologies occasionally produce false-positive and false-negative classifications driven by the underlying structure of the data modality. False positives frequently arise because technical noise and sequencing dropouts can randomly cluster in spatial dimensions. Global statistical metrics routinely misinterpret these artificial clusters as genuine biological signals across all standard modeling approaches. Conversely, false negatives occur when a true biological pattern is restricted to a narrow anatomical layer, such as those found in the mouse cerebellum. In these instances, the large proportion of zero counts across the rest of the tissue mathematically dilutes the localized signal and causes it to fall below standard detection thresholds. As a result, the engineered features of the agent may occasionally favor broad trends over highly concentrated patterns. This susceptibility to artifacts driven by sparsity is a widely documented limitation across the entire field, where high dropout rates simultaneously inflate false positives and obscure localized genes regardless of the detection framework utilized [50].

### 3.5 Deep adaption of pretrained foundation models

We tasked AIDO.Builder with predicting binary DNA–RNA interactions from sequence data alone. The dataset was curated from RNAInter (RNA Interactome Database) [51] and comprised a training, validation, and test set of more than 200K data samples, each consisting of an RNA sequence, a DNA sequence, and a binary interaction label. The agent was given access to pre-trained biological sequence models (AIDO.RNA-650M [36] and AIDO.DNA-300M [37]), but received no guidance on model architecture, training strategy, or hyperparameters.

The agent began by studying the available pre-trained foundation models—reading the provided AIDO.Builder model code to understand loading procedures, tokenization protocols, and embedding extraction. It identified that both AIDO.RNA-650M and AIDO.DNA-300M are RNABert-family [52] encoders sharing the same tokenizer framework, but producing embeddings of different dimensionalities (1,280 for the RNA model, 1,024 for the DNA model). Based on its understanding of the foundation models, the agent designed a dual-encoder architecture in which the two pre-trained models process RNA and DNA sequences independently, their outputs are fused, and a classification head produces a binary interaction prediction (Fig. 5 first panel). The initial design froze both encoder backbones entirely and applied low-rank adaptation [53] (LoRA) to the query and value projections of both encoders’ attention layers, enabling parameter-efficient fine-tuning with only 1.5% of parameters trainable. Mean pooling over non-special tokens was used to obtain fixed-length sequence representations from the variable-length DNA/RNA sequence embedding. The agent implemented a simple fusion strategy by concatenating the DNA and RNA representations followed by a two-layer MLP to predict the class.

**Fig. 5.**
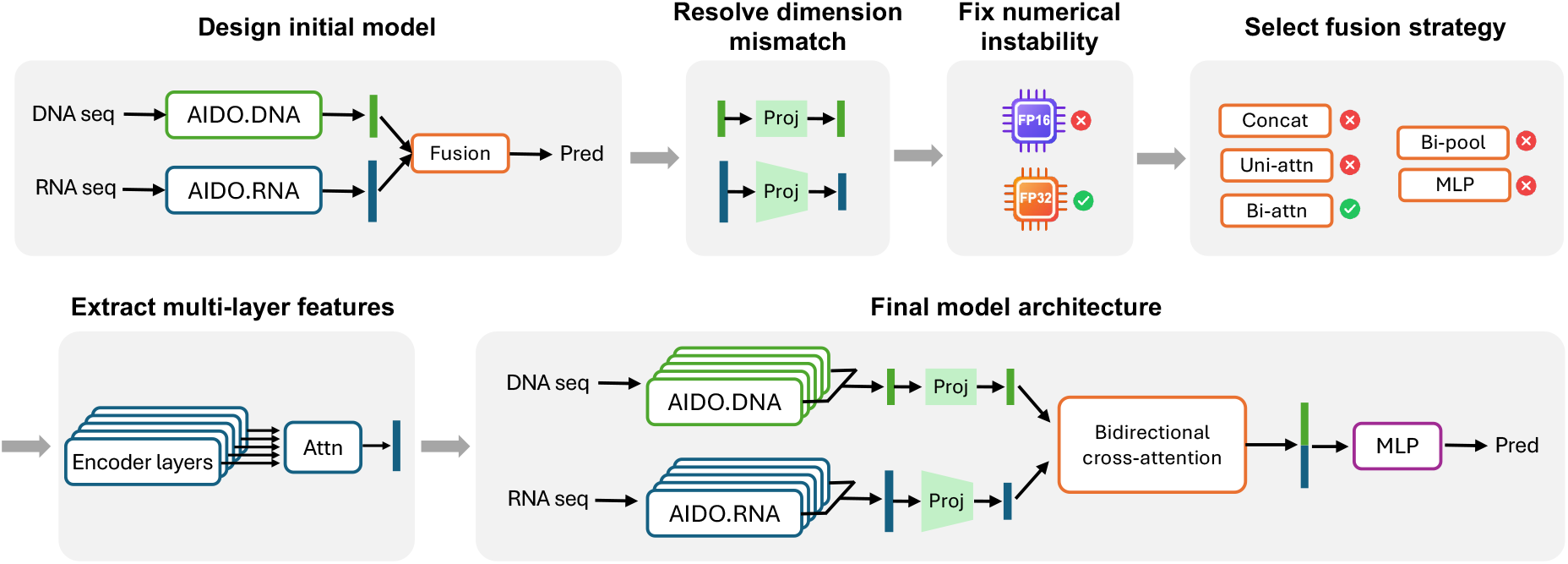
Autonomous design and refinement of a dual-encoder architecture for DNA–RNA interaction prediction. The AIDO.Builder was tasked with predicting binary DNA–RNA interactions from sequence data using pre-trained foundation models (AIDO.RNA-650M and AIDO.DNA-300M) without human guidance. **Top** Left, initial model design: independent encoding of RNA and DNA sequences followed by simple fusion and classification. Middle, iterative debugging: the agent resolved embedding dimensionality mismatch by introducing learned projection modules and stabilized training by replacing FP16 with FP32 precision and gradient clipping. Right, systematic fusion strategy search: among multiple candidates, bidirectional cross-attention emerged as the most effective mechanism for modeling inter-molecular interactions. **Bottom** Left, multi-layer feature extraction: the agent extracted features from the last six encoder layers and aggregated them via attention pooling, resulting in rich molecular features from the AIDO.DNA/RNA encoder. Right, final architecture: the AIDO.DNA and AIDO.RNA encoders encode the DNA and RNA sequences, respectively. Multi-layer features from the last six encoder layers are aggregated via attention pooling, projected to a shared embedding space, and fused using bidirectional cross-attention, followed by an MLP classifier. This autonomous, multi-stage optimization pipeline led to substantial performance improvements from near-random initialization to strong predictive accuracy on the held-out test set.

The initial architecture performed near chance level (validation AUROC 0.53), and the agent undertook a systematic, component-by-component refinement process spanning over 30 steps to diagnose and resolve the underlying issues.

With a numerically stable training pipeline, the agent conducted a rapid prototyping experiment testing five fusion strategies—concatenation, unidirectional attention, bidirectional cross-attention, bilinear pooling, and deep MLP fusion—on a small subset with 500 training samples. Bidirectional cross-attention, in which RNA representations attend to DNA and DNA representations attend to RNA via separate 8-head multi-head attention modules with residual connections and layer normalization [54], emerged as the best-performing strategy. The agent adopted this as the fusion module, producing a 2,048-dimensional concatenated output from the two attended representations.

Despite these improvements, performance plateaued at AUROC 0.61 on the validation set. The agent hypothesized that relying solely on the final hidden state of each encoder discarded useful information captured in intermediate layers. It implemented a multi-layer feature extractor that retrieves outputs from the last *N* encoder layers and combines them via a learned attention-pooling mechanism—a query vector attending over layer outputs to produce a weighted combination. The agent systematically compared configurations using 2, 4, 6, and 8 layers on a validation subset for fast prototyping and found that 6 layers provided the best AUROC (0.683), which it adopted as the default. This component allowed the model to leverage the hierarchical representations learned across multiple depths of the pre-trained encoders. Through systematic diagnosis of the model training, the agent introduced a class-weighted binary cross-entropy loss and a weighted random data sampler for balanced mini-batch construction, directly addressing the false-negative bias.

The final model integrates the cumulative refinements into a cohesive architecture (Fig. 5). Two frozen pre-trained encoders (AIDO.RNA-650M and AIDO.DNA-300M) with LoRA adapters extract multi-layer features from the last 6 encoder layers, which are combined via learned attention pooling across layers within each encoder. The DNA/RNA features are projected to a shared 1,024-dimensional space, where bidirectional cross-attention fusion captures inter-molecular interactions. A three-layer MLP head with layer normalization, ReLU, and dropout produces the final prediction.

On the full held-out test set (48,911 samples), the final model achieved an AUROC of 0.792 and AUPRC of 0.679, with accuracy of 61.8%, precision of 39.0%, recall of 81.4%, and F1 of 52.7% at a threshold of 0.5. Performance improved monotonically through the agent’s iterative refinements: from near-random (AUROC 0.50) with the initial architecture, to 0.683 after multi-layer feature extraction, and finally to 0.792 after the redesigned training strategy.

### 3.6 Iterative refinement of pretrained foundation models

For biological tasks that depend heavily on contextual information, large pretrained models often need architecture and optimization strategies to be refined in response to empirical feedback. We evaluated whether the agent could autonomously execute this iterative refinement process (Fig. 6) using the dipeptidyl peptidase IV (DPP-IV) inhibitory peptide prediction task [55, 56]. This task frames the identification of diabetes therapeutics as a supervised binary classification problem, in which the model predicts bioactivity from a primary amino acid sequence. Given a natural language specification to fine-tune a protein foundation model, the agent first constructed a reproducible baseline pipeline. It implemented data preprocessing and instantiated a small baseline architecture comprising a 35-million-parameter protein language model (ESM2 [57]) paired with a linear classification head. After resolving initial code execution errors, the agent established a functioning reference implementation that achieved an initial AUROC of 0.925 on the held-out test set.

**Fig. 6.**
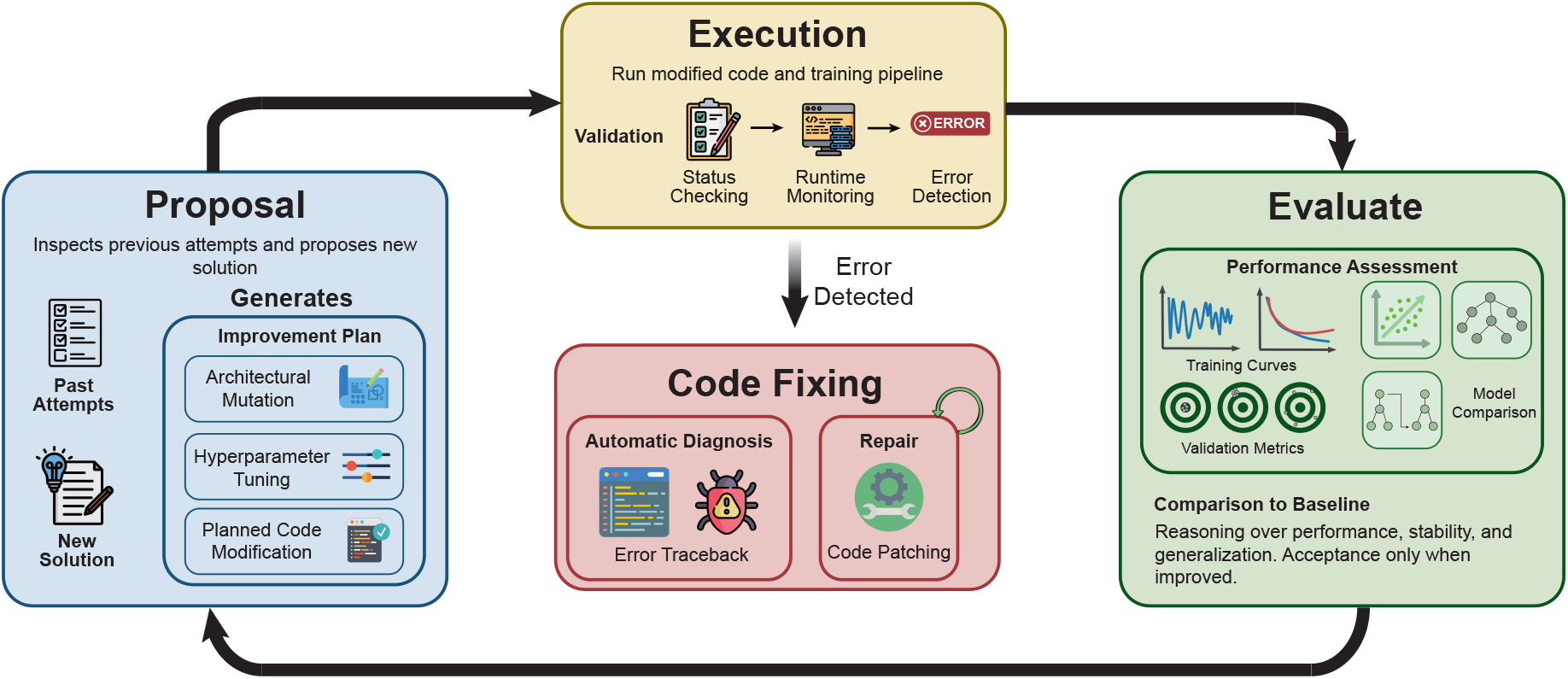
Iterative refinement module for autonomous model improvement. AIDO.Builder refines a baseline training pipeline through a closed loop of proposal, execution, evaluation and code fixing. Guided by previous attempts, the proposal stage generates a new solution through architectural modification, hyperparameter tuning and planned code changes. The modified pipeline is then executed under validation, runtime monitoring and error detection. Successful runs are evaluated using training behaviour, validation metrics, model comparisons and performance relative to baseline, and are retained only when they improve performance, stability or generalization. Failed runs trigger automatic diagnosis and code patching before re-entry into the refinement loop.

Starting from this reference baseline, the agent executed a bounded refinement loop consisting of three iterations (Fig. 7). In each iteration, it analyzed training curves, loss dynamics, and evaluation metrics to propose targeted updates. In the first iteration, the agent focused on the classification head architecture. Analyzing the baseline loss dynamics, it replaced the single linear layer with a multilayer perceptron incorporating residual connections and layer normalization. It also implemented attention pooling to dynamically weight the sequence residues most predictive of DPP-IV inhibition. Noting the absence of overfitting in the baseline training curves, the agent decreased the dropout rate from 0.3 to 0.1. This architectural modification increased the test AUROC from 0.925 to 0.9301.

**Fig. 7.**
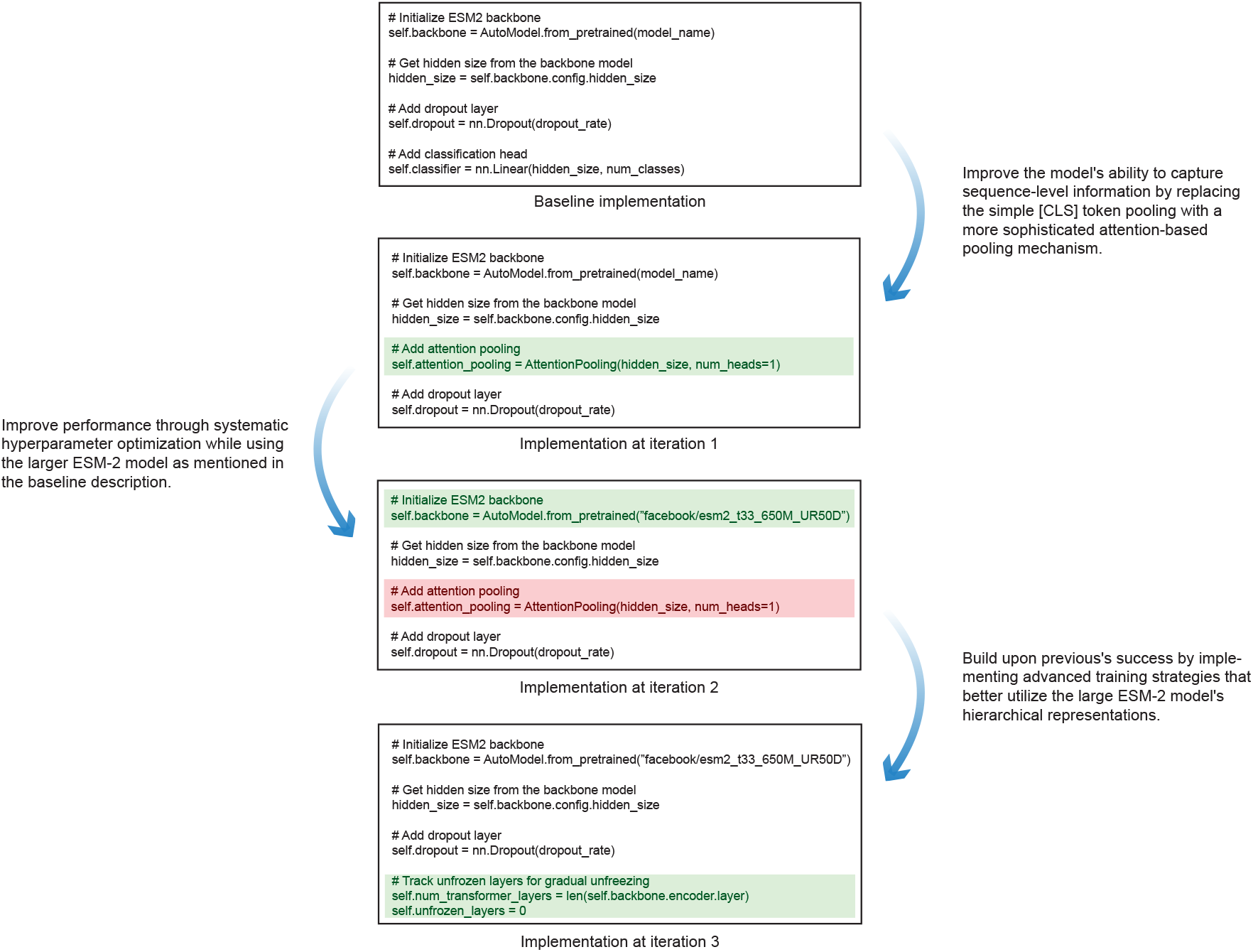
Example of iterative refinement during autonomous adaptation of a protein foundation model. Starting from a reproducible baseline for DPP-IV inhibitory peptide discovery, AIDO.Builder performs three rounds of feedback-guided refinement. In the first iteration, it improves the sequence classifier by redesigning the prediction head and introducing attention-based pooling. In the second iteration, it scales the backbone and restructures optimization to support larger-model training. In the third iteration, it refines fine-tuning dynamics through layer-wise adaptation and stabilized scheduling. Together, these iterations illustrate how empirical training feedback is translated into successive improvements in model architecture, scale and optimization strategy.

**Fig. 8.**
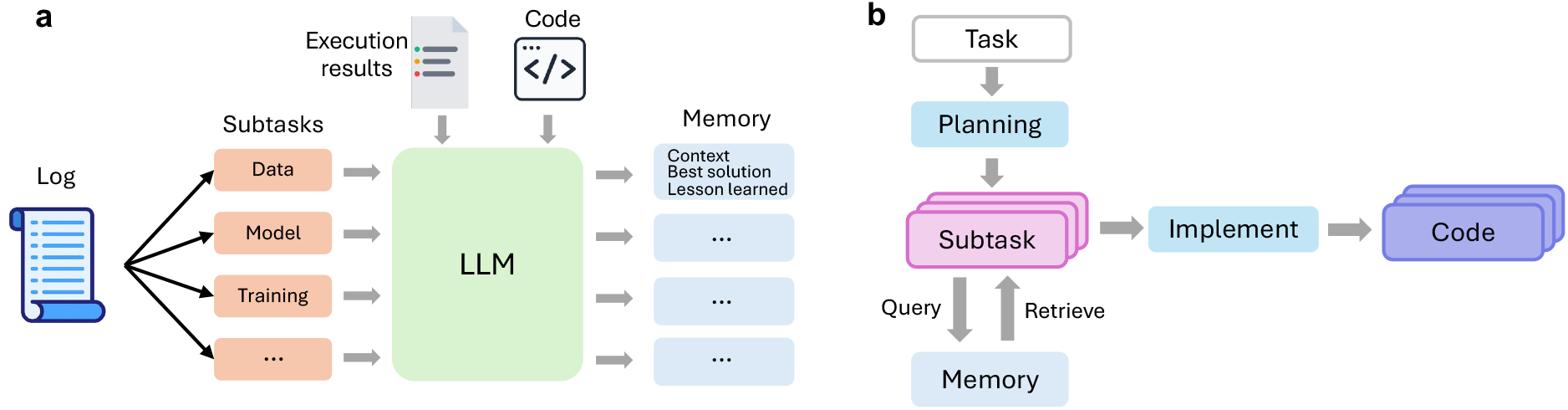
Experience memory for cross-task transfer in the autonomous agent. **a**, Memory creation by post hoc reflection over execution trajectories. After task completion, the agent analyses logs, generated code and execution results, decomposes the trajectory into semantically meaningful subtasks, and converts each into a structured memory record containing the task description, broader context, best solution and lessons learned. These records are embedded and stored in a persistent memory database shared across runs. **b**, Memory retrieval during task execution. For each newly planned subtask, the agent queries the memory store using semantic similarity over task-description embeddings, retrieves relevant prior experiences, and injects them into the implementation prompt to guide code generation. This design enables reuse of successful pipeline patterns, debugging strategies and operational knowledge across biologically distinct tasks without manual curation.

In the second iteration, the agent scaled the core representation by replacing the 35-million-parameter backbone with a 650-million-parameter model. To accommodate the increased memory requirements and stabilize the optimization of this larger model, the agent autonomously restructured the training parameters. It halved the batch size from 16 to 8 to prevent out-of-memory errors and implemented a two-step gradient accumulation process to maintain the effective batch size. Additionally, it reduced the learning rate, increased the dropout rate back to 0.2, increased weight decay for stronger regularization, and extended the early-stopping patience. This coordinated adjustment of model scale and memory management increased the test AUROC to 0.9545.

In the final iteration, the agent shifted its focus to the fine-tuning dynamics of the transformer architecture to prevent the loss of pretrained representations. It introduced a layer-wise learning rate decay, assigning smaller gradient updates to the lower transformer blocks while allowing larger updates near the classification head. It combined this with a gradual, top-to-bottom unfreezing schedule. To further stabilize the training process, the agent applied label smoothing, layer-specific dropout scheduling, and a cosine annealing learning rate schedule with warm restarts. These fine-tuning adjustments yielded a final test AUROC of 0.9656. These results demonstrate that the agent can progressively optimize both model architecture and training dynamics by directly backpropagating empirical execution feedback into structural pipeline revisions.

### 3.7 Agent experience accumulation and memory retrieval

A central question in deploying LLM agents across serial biomedical tasks is whether experience gained on one task can transfer to subsequent, biologically distinct tasks. To investigate this, we examined the agent’s embedding-indexed experience memory (Methods), which accumulates structured records of past solutions and lessons learned and retrieves them via semantic similarity search when the agent begins an implementation step of a new subtask (Fig. 8). The BioDiscoveryBench benchmark [34] is an ideal dataset to demonstrate the agent’s ability to transfer past experience when solving related biomedical tasks. The benchmark requires an agent to iteratively propose genes for perturbation across multiple rounds of a simulated CRISPR screen. It consists of 6 tasks spanning diverse biological contexts (Fig. 9a). The agent was given access to a pool of candidate genes, genomic features, AIDO. Builder foundation models, and observations from each experimental round, and was tasked with maximizing the discovery of hit genes within a fixed budget of five rounds and N (N=128 for all tasks except Scharenberg where N=32) genes in each round. We evaluated the AIDO.Builder on the six CRISPR screen tasks in a sequential order, where the agent can save its experience of solving prior tasks into the memory database and retrieve related memory items when a new task in a different biological context.

**Fig. 9.**
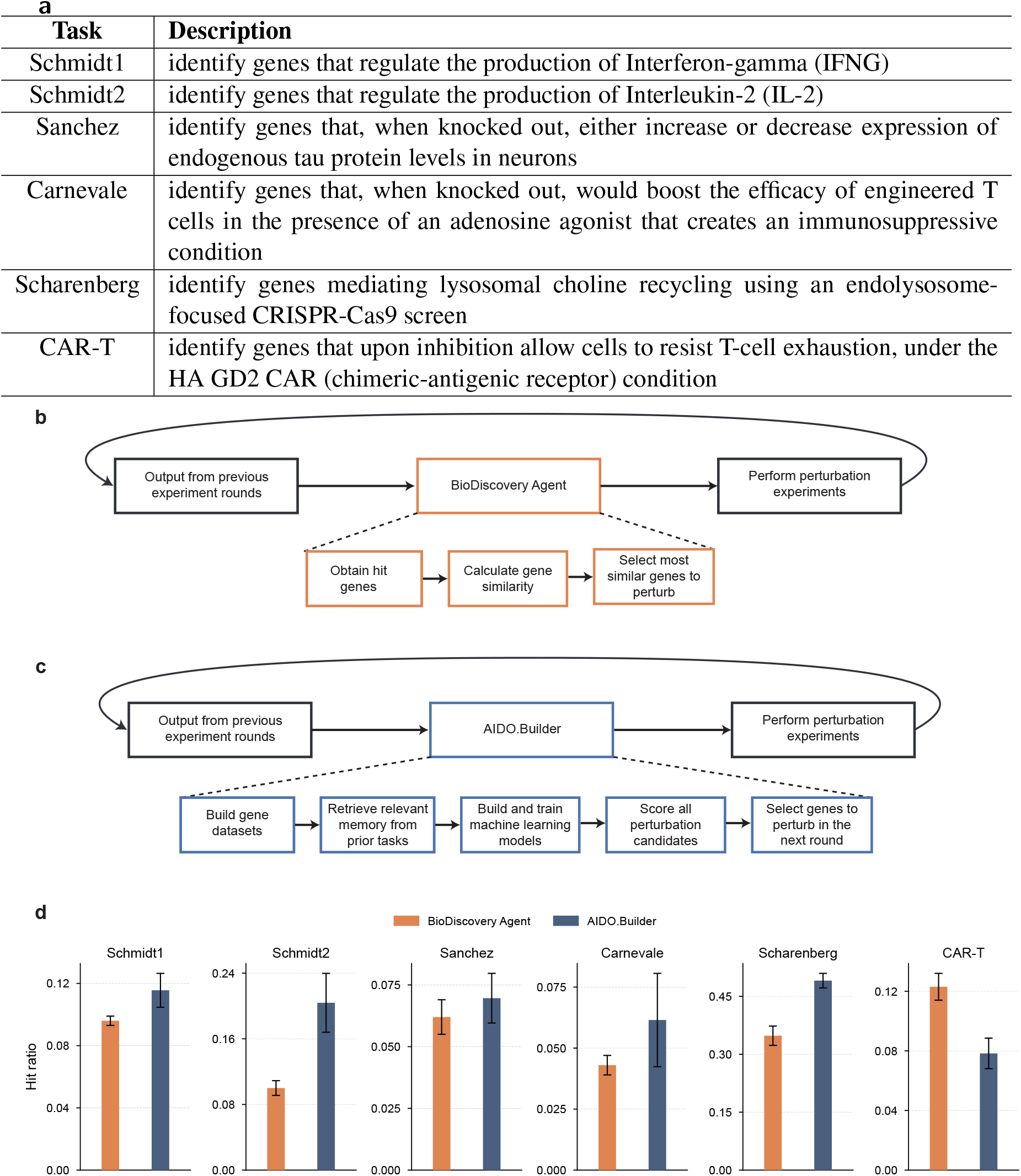
Cross-task transfer of agent experience improves iterative gene discovery in BioDiscoveryBench. **a**, Six BioDiscoveryBench CRISPR-screen tasks used to evaluate cross-task transfer, spanning distinct biological objectives: regulation of interferon-*γ* (Schmidt1), interleukin-2 (Schmidt2), neuronal tau levels (Sanchez), adenosine-resistant engineered T-cell activity (Carnevale), lysosomal choline recycling (Scharenberg) and resistance to CAR-T-cell exhaustion (CAR-T). **b**, Baseline BioDiscovery Agent workflow. The agent iteratively proposes perturbations using outputs from previous rounds together with predefined gene-search tools, including hit-gene identification, gene-similarity calculation and selection of related genes for subsequent perturbation experiments. **c**, AIDO.Builder workflow. After an initial knowledge-driven round, the agent autonomously builds task-specific gene datasets, trains machine-learning models, scores all candidate perturbations and selects genes for the next experimental round; experiences from earlier tasks are stored in persistent memory and retrieved to guide later implementations. **d**, Performance across the six tasks, measured as hit ratio over the fixed experimental budget (mean ± s.d.). AIDO.Builder outperformed BioDiscovery Agent on five of six tasks, with the largest gain on Scharenberg, and also improved performance on Schmidt1, Schmidt2, Sanchez and Carnevale; CAR-T was the only task in which BioDiscovery Agent performed better. These results show that experience accumulated on earlier screens can transfer across biologically distinct tasks, enabling reuse of machine-learning pipeline designs and operational strategies in sequential tasking.

We ran the agent in three independent runs per task, ensuring the memory of a task is used in solving the same task. Performance was measured by hit ratio, defined as the fraction of all possible hit genes in the dataset that the agent successfully identified within its experimental budget (Methods). We compared our AIDO.Builder against the BioDiscovery Agent [34] which proposed the original BioDiscoveryBench. The BioDiscovery Agent was given a tool to search for top 10 genes based on different criteria, such as similar/dissimilar/correlated genes, genes in common tissues, and KEGG enrichment analysis (Fig. 9b).

The AIDO.Builder autonomously constructed a two-phase experimental pipeline for each task. In the first round, it employed pure biological domain knowledge from the base LLM to propose genes to knockout. For subsequent rounds (2-5), the AIDO.Builder built a machine learning model to prioritize untested genes. Due to the small size of the observed perturbation results (with N data points in the first round), the AIDO.Builder decided to use the features from the AIDO.Cell foundation model instead of fine-tuning the foundation model, as the later can easily lead to overfitting given the small size of the training data. The AIDO.Builder derived gene representations from up to 384-dimensional feature vectors combining 256-dimensional PCA embeddings of DepMap Achilles CRISPR-knockout dependency profiles and 128-dimensional AIDO.Cell in-silico knockout embeddings. For most tasks, the agent built a bootstrap ensemble of XGBoost [58] regressors on top of the gene features. After each round, the agent used the accumulated gene scores to train a new model and selected genes for the new round based on the model-predicted scores on untested genes.

The performance comparison between the AIDO.Builder and the BioDiscovery Agent is shown in Fig. 9d. The AIDO.Builder achieved its strongest performance on the Scharenberg dataset, discovering approximately half of all hit genes (mean hit ratio = 0.49 ± 0.019 across three runs) while testing only 15.1% of the gene pool, significantly surpassing BioDiscovery Agent’s 0.348 ± 0.025 (*P <* 0.001). On the two Schmidt tasks (IL-2 and IFNG), AIDO.Builder achieved hit ratios 0.116 ± 0.011 and 0.204 ± 0.036, respectively, also clearly outperforming BioDiscovery Agent’s 0.096 ± 0.003 (*P* = 0.086) on Schmidt1 and 0.010 ± 0.009 (*P* = 0.035) on Schmidt2. On the harder tasks of Sanchez and Carnevale, AIDO.Builder’s hit ratio is also higher than BioDiscovery Agent (0.070 ± 0.010 vs. 0.062 ± 0.007 on on Sanchez (*P* = 0.31); 0.062 ± 0.019 vs. 0.043 ± 0.004 on Carnevale (*P* = 0.23)). The CAR-T task is the only task where BioDiscovery Agent’s hit ratio of 0.123 ± 0.009 outperformed AIDO.Builder’s 0.0783 ± 0.010 (*P* = 0.0065). These results demonstrated AIDO.Builder’s ML model building is generally more effective than BioDiscovery Agent’s gene search method.

We performed a detailed analysis of the AIDO.Builder’s ability to use memory items accumulated in prior BioDiscovery tasks. We found that cross-task retrieval—where experiences from a biologically distinct screen informed the current task—was pervasive and beneficial. Cross-task memory transfer operated through two principal channels. First, and most impactful, was the transfer of ML pipeline architecture. The adaptive XGBoost bootstrap ensemble was first developed during early Schmidt1 IFNG screen runs and subsequently propagated via memory to every other task. When the agent began building an ML model for the Scharenberg lysosomal choline recycling screen, it retrieved experiences from prior Schmidt1 IFNG experiments that specified an effective architecture: “K=3 bootstrap ensemble, adaptive hyperparameters scaled by training set size, UCB acquisition with decaying alpha, quality gate using 5-fold CV Spearman.” The agent adopted this template wholesale, achieving cross-validated Spearman correlations of 0.22-0.43 on Scharenberg from the first ML round onward. Similarly, when building an ML pipeline for the CAR-T screen, the agent retrieved the Schmidt2 IL-2 experiment’s ML pipeline, in particular its quality gate mechanism, where it estimated model reliability via 5-fold cross-validated Spearman correlation on the training data, and when this metric fell below a threshold, the agent fell back to a heuristic strategy for genes selection. This architectural transfer effectively eliminated the need to re-derive the active learning strategy from scratch for each new biological screen.

Second, cross-task memory transferred critical operational patterns that prevented common failure modes. A recurring example was the environment variable timing lesson: the agent learned from an early Carnevale experiment that the BIODISCOVERY_EXPERIMENT_ID and DATASET_NAME variables must be set before importing the BioDiscovery Python module. This lesson, stored in memory, was retrieved and applied in all subsequent tasks across all runs, preventing a class of initialization failures. Another widely transferred operational pattern was the over-submission strategy: a Schmidt1 experiment discovered that submitting approximately 2x the round budget ensured full utilization even when some gene names were invalid or already tested, and this was subsequently retrieved and adopted by the other tasks.

## 4 Discussion

The automation of machine learning model development represents a critical step toward scaling data-driven biomedical discovery. In this study, we demonstrated that an autonomous agentic system, AIDO.Builder, can successfully navigate the end-to-end lifecycle of predictive modeling across diverse biological modalities. Rather than functioning merely as a code-generation assistant, the agent operates as an independent experimentalist that formulates domain-appropriate architectures, resolves runtime integration errors, aligns validation strategies with target metrics, and iteratively refines training dynamics based on empirical feedback. By successfully synthesizing both task-specific architectures from scratch and fine-tuning large-scale foundation models, the system illustrates a transition from manual trial-and-error engineering to systematic, automated pipeline construction.

A primary bottleneck in biomedical data science is the fragility of computational pipelines, where minor data formatting anomalies or dependency conflicts frequently derail model development. The agent’s capacity for systematic error resolution addresses this friction directly. As observed in the small-molecule and protein fitness tasks, the agent’s ability to autonomously parse tracebacks, substitute missing dependencies, and adjust tensor dimensions ensures that pipeline execution does not terminally halt at the first software exception. Furthermore, the independent trials on the Q8EG35 dataset highlight a fundamental operational principle for autonomous systems, demonstrating that protocol stability and defensive error handling often dominate architectural complexity. The agent’s capacity to execute fallback pathways when fundamental representation assumptions fail demonstrates a necessary failure-containment mechanism, ensuring protocol adherence even under suboptimal modeling conditions.

Beyond execution robustness, AIDO.Builder exhibits domain-aware optimization that mirrors the empirical refinement process of human researchers. In spatial transcriptomics, the agent moved beyond generic global statistics to engineer localized spatial graphs and autonomously implemented sanity checks to mitigate data leakage. In the DPP-IV foundation model task, it successfully scaled from a 35-million-parameter to a 650-million-parameter backbone by dynamically adjusting batch sizes, gradient accumulation, and layer-wise learning rates in response to training loss and memory constraints. These behaviors indicate that large language models, when embedded in reproducible execution environments, can effectively backpropagate empirical training signals into structural pipeline revisions.

Building upon these isolated model optimizations, AIDO.Builder further demonstrates the capacity to architect complex, multimodal systems and accumulate scientific experience over time. In the DNA-RNA interaction task, the agent autonomously engineered a dual-encoder architecture, resolving dimensionality mismatches and systematically evaluating fusion strategies to converge on bidirectional cross-attention without human guidance. Crucially, the system extends this autonomous problem-solving into cumulative learning via a retrieval-augmented memory module. As demonstrated in the BioDiscoveryBench CRISPR screens, the agent successfully transferred architectural templates, operational safeguards, and dataset-specific modeling insights across distinct biological domains. This cross-task memory transfer mimics the accumulated knowledge of a human research lab, enabling the agent to progressively improve its efficiency on novel tasks without re-deriving complex optimization strategies from scratch.

Despite these capabilities, fully autonomous model development remains constrained by several fundamental factors. First, the efficacy of the agentic system is fundamentally restricted by the reasoning, planning, and coding capabilities of the underlying large language model [32]. Because the agent relies on the base model to interpret complex tracebacks and synthesize architectures, its ceiling for scientific innovation is intrinsically tied to the model’s training distribution and context limits. Second, the current system relies on well-defined evaluation metrics, curated data splits, and explicit task formulations. Applying this agentic framework to entirely unstructured, unannotated raw biological data, where the objective function itself must be discovered, remains an open challenge. Finally, achieving the ultra-long-term, intervention-free autonomous operation necessary to solve such open-ended problems will require more sophisticated mechanisms for self-learning and self-evolution. While our retrieval-augmented memory module enables effective cross-task knowledge transfer and the accumulation of explicit operational patterns, the system cannot yet autonomously invent entirely novel algorithmic paradigms or fundamentally rewrite its own orchestration logic over extended time horizons.

Ultimately, autonomous agentic model building provides a scalable approach for democratizing advanced machine learning in biomedicine. By automating the substantial engineering overhead required to construct, debug, and optimize computational pipelines, systems like AIDO.Builder allow researchers to focus on high-level experimental design, data curation, and hypothesis generation. As underlying large language models continue to improve in long-horizon reasoning and code synthesis, the integration of such autonomous agents into the scientific workflow will accelerate the translation of general algorithmic advances into reliable, task-specific biomedical tools.

